# PICKLES v3: the updated database of pooled in vitro CRISPR knockout library essentiality screens

**DOI:** 10.1101/2022.09.16.508307

**Authors:** Lance C Novak, Juihsuan Chou, Medina Colic, Christopher A. Bristow, Traver Hart

## Abstract

PICKLES (https://pickles.hart-lab.org) is an updated web interface to a freely available database of genome-scale CRISPR knockout fitness screens in human cell lines. Using a completely rewritten interface, researchers can explore gene knockout fitness phenotypes across cell lines and tissue types and compare fitness profiles with fitness, expression, or mutation profiles of other genes. The database has been updated to include data from three CRISPR libraries (Avana, Score, and TKOv3), and includes information from 1,162 whole-genome screens probing the knockout fitness phenotype of 18,959 genes. Source code for the interface and the integrated database are available for download.

## Introduction

Loss of function fitness screens in cell lines offer the opportunity for integrative and comparative analyses to identify background-specific genetic vulnerabilities. While the technology behind CRISPR-mediated genetic and epigenetic perturbation has evolved rapidly, by far the largest data sources remain the genome-scale CRISPR/Cas9 knockout screens in hundreds of cancer cell lines. Under the various banners of The Cancer Dependency Map (DepMap) (1), Project Achilles (2), and Project Score (3), as well as additional independent projects (4–10), roughly a thousand cancer and other human cell lines have been screened using whole-genome CRISPR/Cas9 knockout libraries.

New bioinformatics tools have been developed to process the data from this new technology. A number of algorithms are available to derive gene-level fitness scores from screens where multiple reagents target each gene. The Project Achilles algorithm, CERES (2), has been supplanted by CHRONOS (11). Likewise, we have updated our original BAGEL algorithm (12) with an improved BAGEL2 (13), a variation of which is used in the Project Score release (3). We have additionally developed a Z-score based algorithm, based on a Gaussian mixture model of the fold change distribution of CRISPR guide RNAs, that is more sensitive in detecting proliferation suppressor genes – those whose knockout increases cell proliferation rates, the opposite phenotype of essential genes (14).

Here we present an updated PICKLES: the database of Pooled In vitro CRISPR Knockout Library Essentiality Screens. PICKLES presents an easy to use interface to CRISPR screen data from three commonly used knockout libraries, enabling simple visualization of gene essentiality scores across cell line and tumor type and comparison of knockout fitness profiles with fitness or gene expression profiles of other genes.

### Data sources and preprocessing

In this PICKLES update we include knockout fitness data from three CRISPR libraries (Table 1), used in Project Achilles (Avana) and Project Score (Score) as well as several smaller independent screening projects that screened multiple cell lines (4–10) using the TKOv3 library (15) (Figure 1). We acquired raw read count data and processed each dataset with the BAGEL2 (13) pipeline, where essential genes have positive logBF scores. We additionally processed the same data with the Z-score pipeline from (14), where negative scores indicate loss of fitness and positive scores indicate faster cell proliferation on gene knockout (e.g. tumor suppressors). We additionally downloaded the processed CHRONOS (11) scores from DepMap portal for comparison here. Cell line metadata was drawn from DepMap or manually annotated.

**Table 1.**
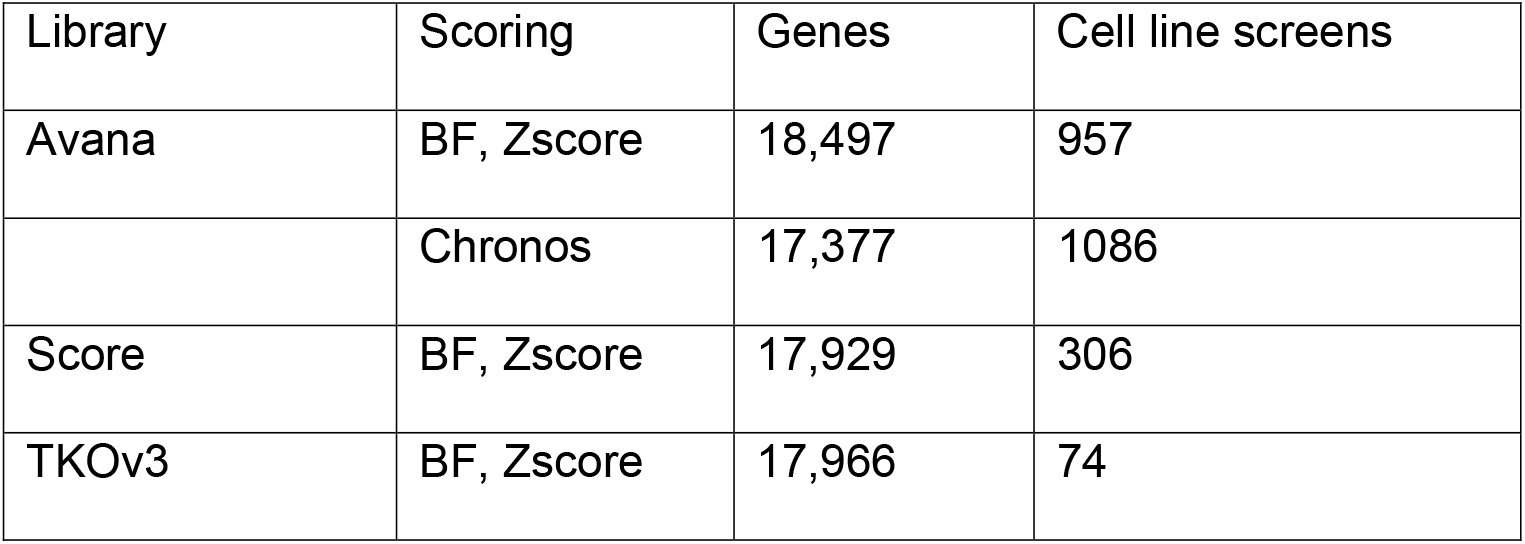
Data sources used.

**Figure 1:**
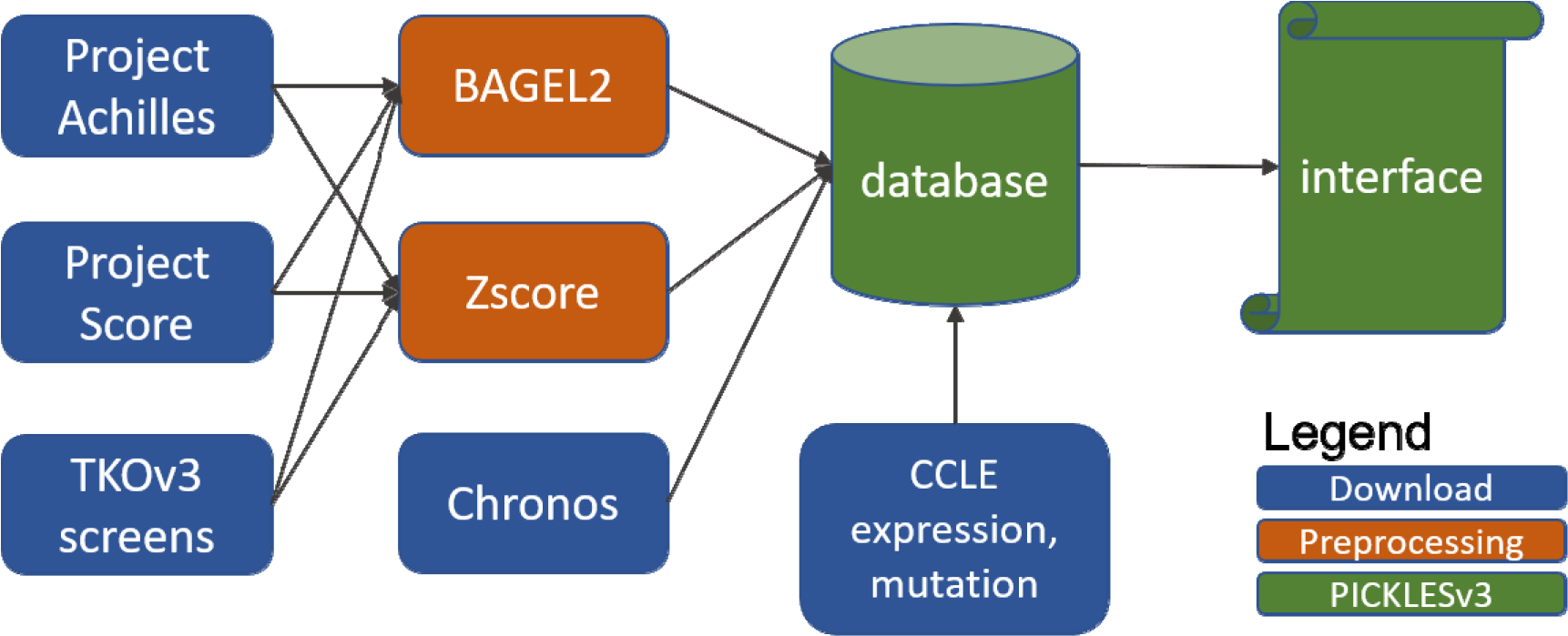
Data preprocessing pipeline. Read count data from the Project Achilles and Project Score, as well as publications with several TKOv3 screens, are downloaded and gene fitness scores are calculated with BAGEL2 and Zscores. Gene fitness scores from Chronos and CCLE molecular profile data are downloaded and integrated with other data in the PICKLES database.

Processed gene expression data, in log2(TPM) format, was downloaded from the CCLE portal (16).

### Database interface and tutorial

The index page of the PICKLES v3 website offers the user the choice of dataset and essentiality scoring scheme to display (Figure 2). Upon selecting a query gene from the text box, the cell lines corresponding to the selected data set are ranked by the chosen scoring metric and plotted in the display panel. The query gene’s essentiality score in each cell line is represented as a point on the graph, color coded by the linage of the cell line. While no thresholding is displayed, reasonable thresholds for an essential gene are BF > 10, Z-score < −4, or Chronos score < −0.75, while a Z-score > 4 typically indicates a proliferation suppressor gene. The “Cancer types” tab displays the same data as the summary page, but with a swarm plot grouped by lineage, with lineages ordered by median essentiality score.

**Figure 2.**
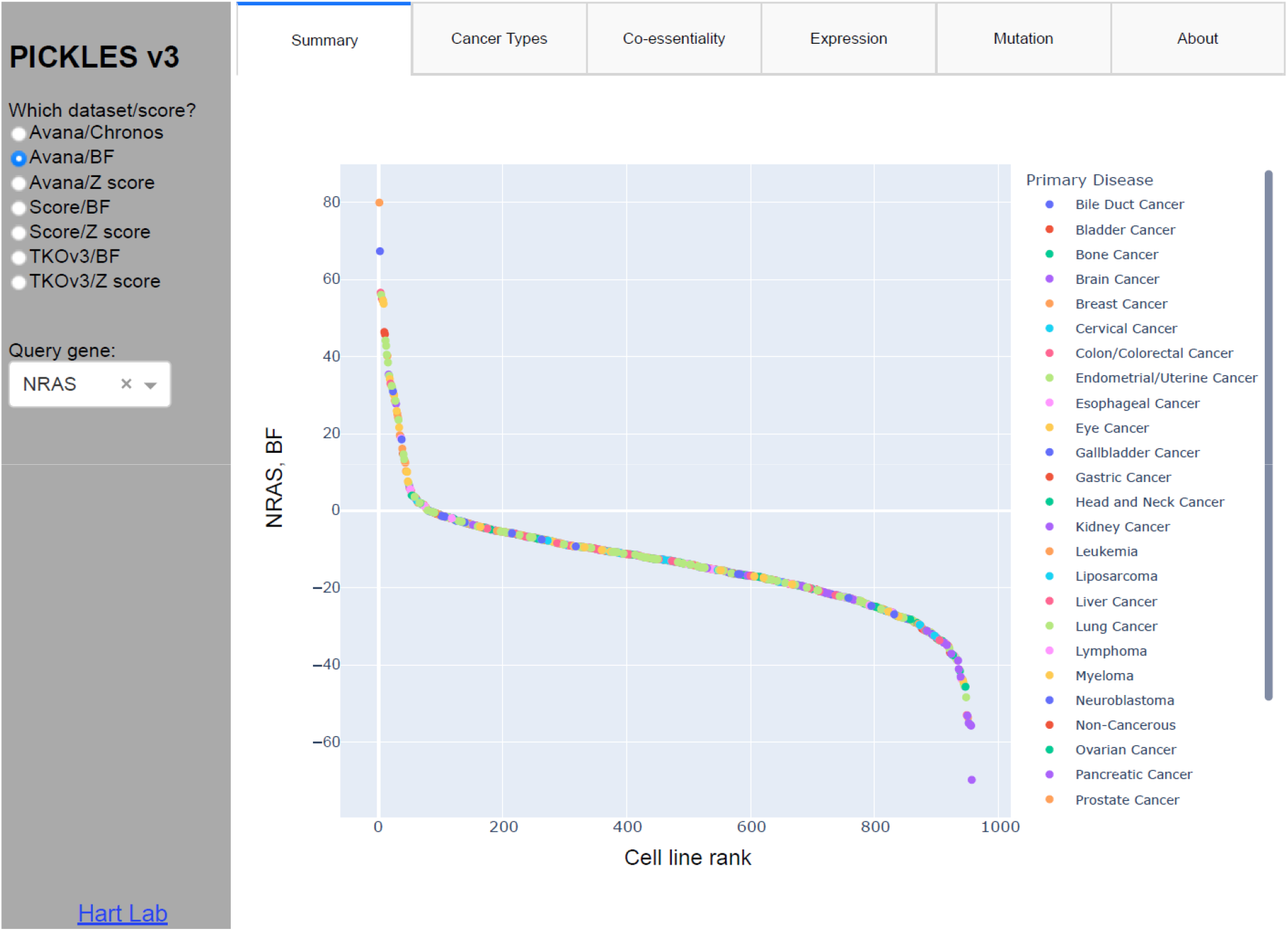
PICKLES v3 summary view. User chooses the dataset and the gene of interest. Ranked scatterplot across relevant cell lines is dynamically generated in the display window. Interactive functions include zoom, pan, select, mouseover for detail, and download image as PNG.

The PICKES v3 interface is implemented in Python/Dash using the Plotly(17) library and includes the native functionality of the Plotly system. On mouseover, each point will display the cell line name, its lineage, and the essentiality score. A single click of the primary disease (legend at right; Figure 2) will remove that lineage from the plot, and another single click returns it. A double click removes all points except the chosen lineage. In addition, hovering the mouse over the plot brings a context menu where the user can zoom, pan, and autoscale the view, and save the modified image in PNG format.

A number of prior works have demonstrated that genes with correlated knockout fitness profiles tend to operate in the same biological process or pathway (they are “co-functional”) (18–23). The coessentiality tab offers the ability to compare The Co-essentiality tab offers the ability to display and measure the essentiality score of the query gene against a second, “comparison” gene in the same dataset, and in addition calculates an ordinary least squares fit between the two profiles, the coefficients of which can be viewed by mouseover.

The coessentiality tab also displays the top ten positive and negative correlates in the data set. A recent study by Wainberg et al (23) indicated that the combination of covariance “whitening” followed by ordinary least squares (OLS) offered significantly improved performance over Pearson correlation in detecting co-functional relationships from coessentiality data. A subsequent study by Gheorghe et al (24) confirmed the performance boost from whitening but showed that, after whitening, Pearson correlation and OLS are mathematically identical. We therefore apply the Cholesky whitening as described in Wainberg et al before determining the top and bottom ranked correlates by PCC. Figure 3B shows the coessentiality of query gene NRAS and comparison gene SHOC2, an important effector of the MAP kinase pathway and the top correlate of NRAS.

**Figure 3.**
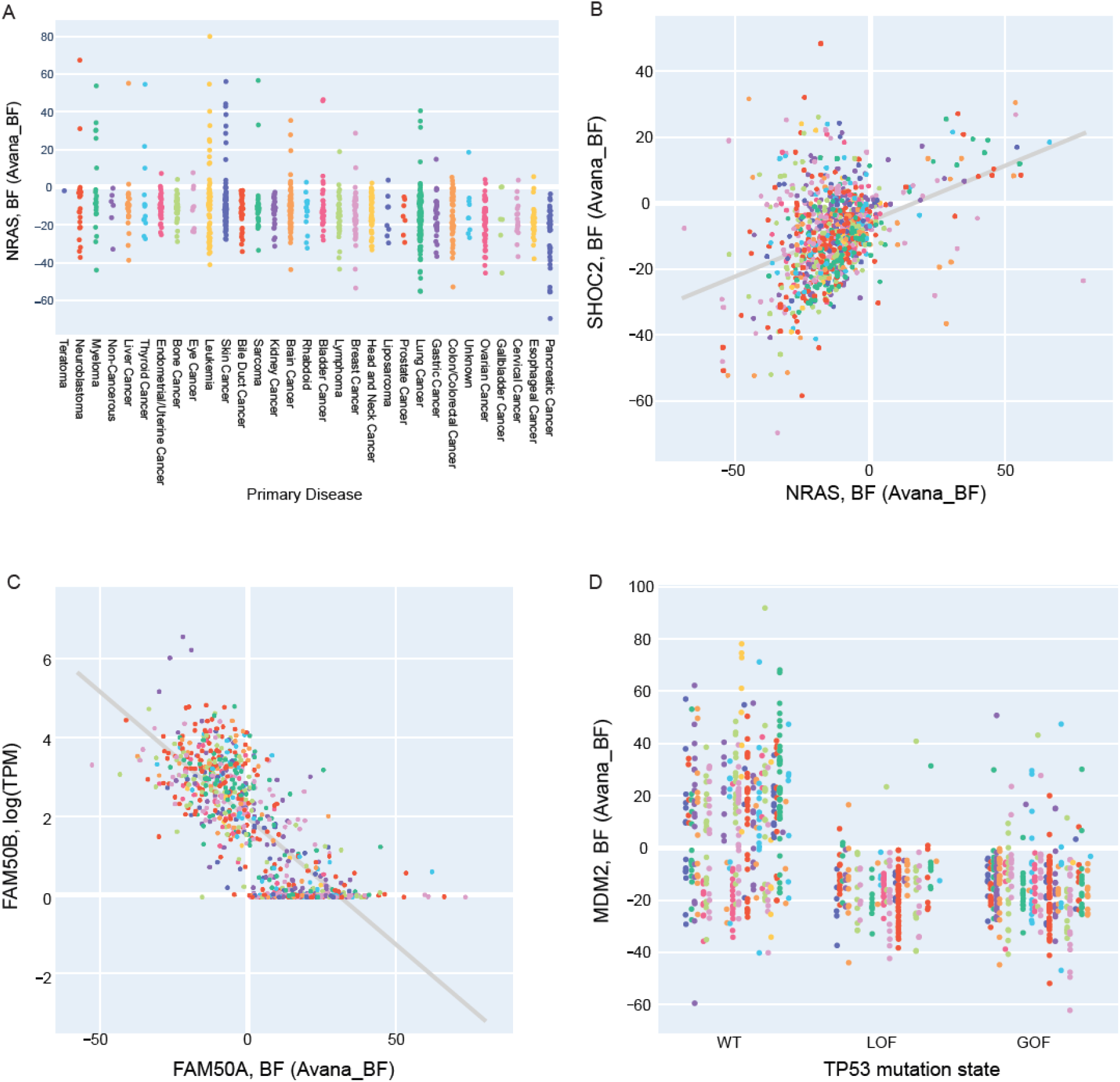
Display options for PICKLES v3 data. (a) Display by cancer types (query gene: NRAS; data source Avana/BF). (B) Co-essentiality (query gene: NRAS; comparison gene: SHOC2). (c) Essentiality vs. Expression (query gene: FAM50A; comparison gene FAM50B). (d) Essentiality vs. Mutation (query gene: MDM2; comparison gene TP53) (e) About and download.

Under the expression tab, the user can view the query gene’s essentiality profile vs. the comparison gene’s gene expression profile in the same cells. Setting the query and comparison genes to the same value allows the user to identify essential genes with tissue-specific expression profiles (e.g. melanocyte transcription factor MITF), while comparing different genes can generate or confirm hypotheses about paralog synthetic lethality. For example, largely uncharacterized paralogs FAM50A and FAM50B are synthetic lethal (25, 26) in digenic knockout screens, and absence of FAM50B gene expression is associated with dependence on FAM50A in cells (Figure 3C).

The mutations tab shows the distribution of essentiality scores in the presence or absence of a LOF or GOF mutation in the comparison gene, separated by lineage/disease type. Mutation data is inferred from CCLE genotyping data available at (16). Briefly, for each gene in each cell line, if a mutation is predicted deleterious, it is classified as Loss of Function (LOF); a deleterious mutation at a hotspot is classified as a Gain of Function (GOF). Figure 3D provides a visualization the relationship between TP53 mutation and the essentiality of MDM2, a key regulator of wildtype p53 protein.

Finally, the about tab offers a brief description of the contents of the PICKLES interface, references, information on how to cite the database, and a link to the PICKLES master data file.

### Implementation

The web interface is implemented in Python/Dash, which uses the Plotly library to dynamically generate interactive graphics for web presentation. The entire application resides in a single ~500-line Python executable, downloadable from our github repository at https://github.com/hart-lab/PicklesV3. On startup, the application loads the entire database, a text file with 21.4 million rows, into memory, consuming < 6% of available RAM on a 128GB consumer-class PC.

## Conclusions

This update to the PICKLES database increases the number of cell lines more than 20-fold and, for most cell lines, offers fitness scores from multiple analysis pipelines. Using a simple and responsive interface, researchers can use not only explore the fitness profiles of their genes of interest but also examine how those profiles change across data set and informatics pipeline. Inclusion of gene expression and mutation data also allows analysis of variation relative to mRNA level and the presence of deleterious mutations in specific genes.

## Funding

JC, MC, and TH were supported by NIGMS grant R35GM130119. TH is a CPRIT Scholar in Cancer Research. This work was supported by the Andrew Sabin Family Foundation Fellowship and by NCI Cancer Center Support Grant P30CA16672.

## Conflict of interest statement

None declared.

## Notes

https://pickles.hart-lab.org/

### Competing Interest Statement

The authors have declared no competing interest.

